# Heparan sulfate proteoglycans as attachment factor for SARS-CoV-2

**DOI:** 10.1101/2020.05.10.087288

**Authors:** Lin Liu, Pradeep Chopra, Xiuru Li, Kim M. Bouwman, S. Mark Tompkins, Margreet A. Wolfert, Robert P. de Vries, Geert-Jan Boons

## Abstract

Severe acute respiratory syndrome-related coronavirus 2 (SARS-CoV-2) is causing an unprecedented global pandemic demanding the urgent development of therapeutic strategies. Microarray binding experiments using an extensive heparan sulfate (HS) oligosaccharide library showed that the receptor binding domain (RBD) of the spike of SARS-CoV-2 can bind HS in a length-and sequence-dependent manner. Hexa- and octa-saccharides composed of IdoA2S-GlcNS6S repeating units were identified as optimal ligands. Surface plasma resonance (SPR) showed the SARS-CoV-2 spike protein binds with much higher affinity to heparin (K_D_ = 55 nM) compared to the RBD (K_D_ = 1 μM) alone. We also found that heparin does not interfere in angiotensin-converting enzyme 2 (ACE2) binding or proteolytic processing of the spike. Our data supports a model in which HS functions as the point of initial attachment for SARS-CoV-2 infection. Tissue staining studies using biologically relevant tissues indicate that heparan sulfate proteoglycan (HSPG) is a critical attachment factor for the virus. Collectively, our results highlight the potential of using HS oligosaccharides as a therapeutic agent by inhibiting SARS-CoV-2 binding to target cells.

## INTRODUCTION

The SARS-CoV-2 pandemic demands urgent development of therapeutic strategies. An attractive approach is to interfere in the attachment of the virus to the host cell.^1^ The entry of SARS-CoV-2 into cells is initiated by binding of the transmembrane spike (S) glycoprotein of the virus to angiotensin-converting enzyme 2 (ACE2) of the host.^2^ SARS-CoV is closely related to SARS-CoV-2 and employs the same receptor.^3^ The spike protein of SARS-CoV-2 is comprised of two subunits; S1 is responsible for binding to the host receptor, whereas S2 promotes membrane fusion. The C terminal domain (CTD) of S1 harbors the receptor binding domain (RBD).^4^ It is known that the spike protein of a number of human coronaviruses can bind to a secondary receptor, or co-receptor, to facilitate cell entry. For example, MERS-CoV employs sialic acid as co-receptor along with its main receptor DPP4.^5^ Human CoV-NL63, which also utilizes ACE2 as the receptor, uses heparan sulfate (HS) proteoglycans, as a co-receptor.^6^ It has also been shown that entry of SARS-CoV pseudo-typed virus into Vero E6 and Caco-2 cells can substantially be inhibited by heparin or treatment with heparin lyases, indicating the importance of HS for infectivity.^7^

There are indications that the SARS-CoV-2 spike also interacts with HS. One early report showed that heparin can induce a conformation change in the RBD of SARS-CoV-2.^8^A combined SPR and computational study indicated that glycosaminoglycans can bind to the proteolytic cleavage site of the S1 and S2 protein.^9-10^ Several reports have indicated that heparin or related structures can inhibit the infection process of SARS-CoV-2 in different cell lines.^11-14^

HS are highly complex *O*- and *N*-sulfated polysaccharides that reside as major components on the cell surface and extracellular matrix of all eukaryotic cells.^15^ Various proteins interact with HS thereby regulating many biological and disease processes, including cell adhesion, proliferation, differentiation, and inflammation. They are also used by many viruses, including herpes simplex virus (HSV), Dengue virus, HIV, and various coronaviruses, as receptor or co-receptor.^16-18^

The biosynthesis of HS is highly regulated and the length, degree, and pattern of sulfation of HS can differ substantially between different cell types. The so-called “*HS sulfate code hypothesis”* is based on the notion that the expression of specific HS epitopes by cells makes it possible to recruit specific HS-binding proteins, thereby controlling a multitude of biological processes.^19-20^ In support of this hypothesis, several studies have shown that HS binding proteins exhibit preferences for specific HS oligosaccharide motifs.^21-22^ Therefore, we were compelled to investigate whether the spike of SARS-CoV-2 recognizes specific HS motifs. Such insight is expected to pave the way to develop inhibitors of viral cell binding and entry.

Previously, we prepared an unprecedented library of structurally well-defined heparan sulfate oligosaccharides that differ in chain length, backbone composition and sulfation pattern.^23-24^ This collection of HS oligosaccharides was used to develop a glycan microarray for the systematic analysis of selectivity of HS-binding proteins. Using this microarray platform in conjugation with detailed binding studies, we found that the RBD domain of SARS-CoV-2-spike can bind HS in a length- and sequence-dependent manner, and the observations support a model in which the RBD confers sequence selectivity, and the affinity of binding is enhanced by additional interactions with other HS binding sites in for example the S1/S2 proteolytic cleavage site.^9^ In addition, it was found that heparin does not interfere in ACE binding or proteolytic processing of the spike. Tissue staining studies using biologically relevant tissues indicate that heparan sulfate proteoglycans (HSPG) is a critical attachment factor for the virus.

## RESULTS AND DISCUSSION

Surface plasma resonances (SPR) experiments were performed to probe whether the RBD domain of SARS-CoV-2 spike protein can bind with heparin. Biotinylated heparin was immobilized on a streptavidin-coated sensor chip and binding experiments were carried out by employing as analytes different concentrations of RBD, monomeric spike protein and trimeric spike protein of SARS-CoV-2. The spike glycoprotein of SARS-CoV-2 (S1+S2, extra cellular domain, amino acid residue 1-1213) was expressed in insect cells having a C-terminal His-tag.^25-26^ Recombinant SARS-CoV-2-RBD, containing amino acid residue 319-541, was expressed in HEK293 cells also with a C-terminal His-tag.^25-26^ The spike protein trimer, having the furin cleavage site deleted and bearing with two stabilizing mutations, was expressed in HEK293 cells with a C-terminal His-tag. Representative sensorgrams are shown in **Fig. 1**. K_D_ values were determined using a 1:1 Langmuir binding model.

**Figure 1.**
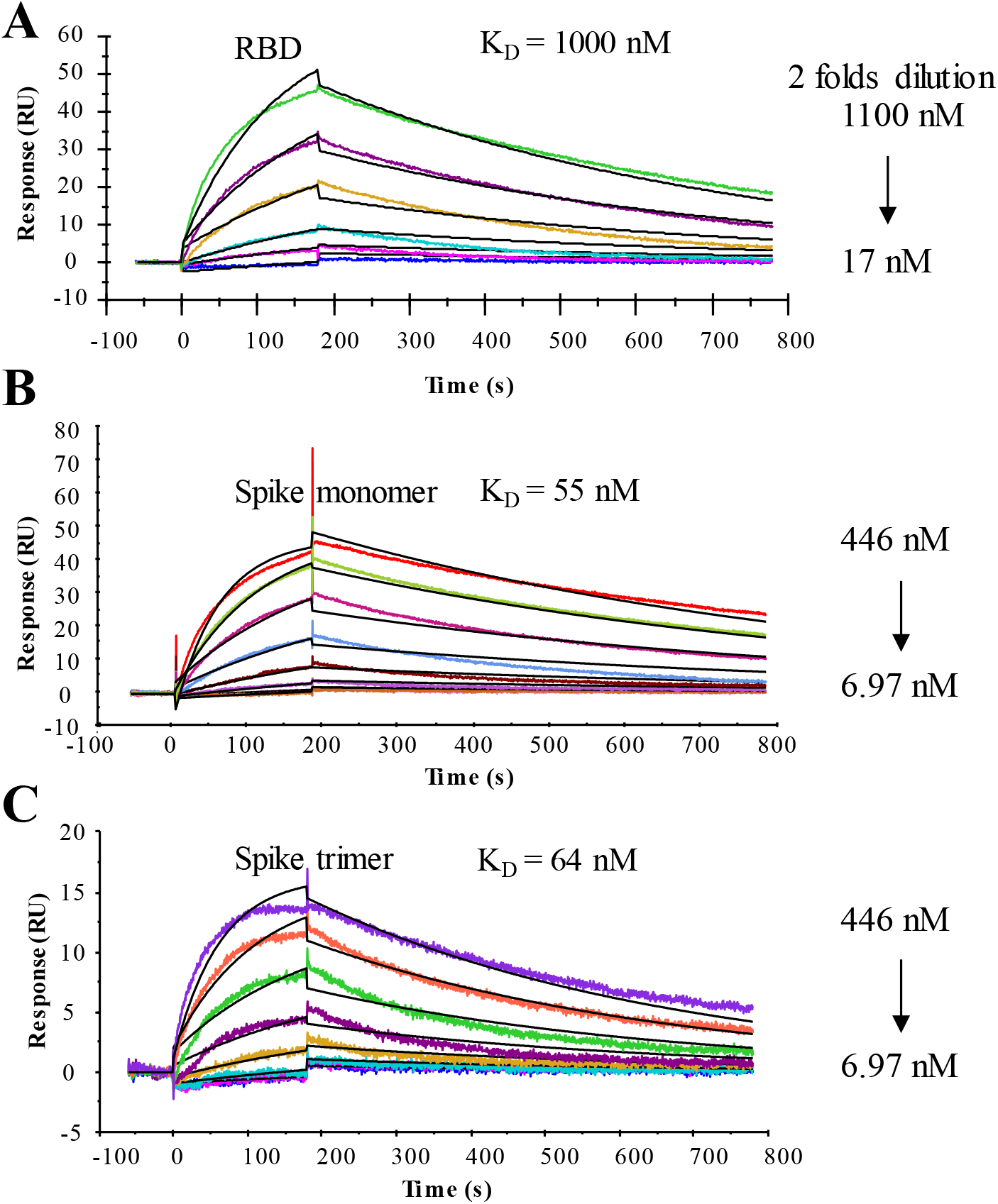
SPR sensorgrams representing the concentration-dependent kinetic analysis of the binding of immobilized heparin with SARS-CoV-2 related proteins (A) RBD, (B) spike monomer, and (C) spike trimer.

The RBD domain binds to heparin with a moderate affinity having a K_D_ value of ∼1 μM. The full-length monomeric spike protein showed a much higher binding affinity with a K_D_ value of 55 nM. Previously reported computational studies have indicated that the RBD domain may harbor an additional HS binding domain located either within or adjacent to the receptor binding motif.^14, 27^ It has also been suggested that another HS-binding site

reside in the S1/S2 proteolytic cleavage site of the spike of the S2 domain.^9^ Thus, the high affinity of the monomeric spike protein probably is due to the presence of additional binding site in the spike protein, which greatly enhanced its binding to heparin. The trimeric spike protein displayed a similar binding affinity (K_D_ = 64 nM) as the monomer. One of the putative heparin binding sites in the trimeric spike protein, the S1/S2 proteolytic cleavage site was mutated.^25^ Thus, a possible increase in avidity due to multivalency may have been off-set by a lack of a secondary binding site. Intrigued by these results, we examined if the SARS-CoV-2 proteins bind to heparan sulfate in a sequence preferred manner. We have developed an HS microarray having well over 100 unique di-, tetra-, hexa-, and octa-saccharides differing in backbone composition and sulfation pattern^23-24^ (**Fig. 2C**). The synthetic HS oligosaccharides contains an anomeric aminopentyl linker allowing printing on *N*-hydroxysuccinimide (NHS)-active glass slides. The HS oligosaccharides were printed at 100 µM concentration in replicates of 6 by non-contact piezoelectric printing. The quality of the HS microarray was validated using various well characterized HS-binding proteins.

**Figure 2.**
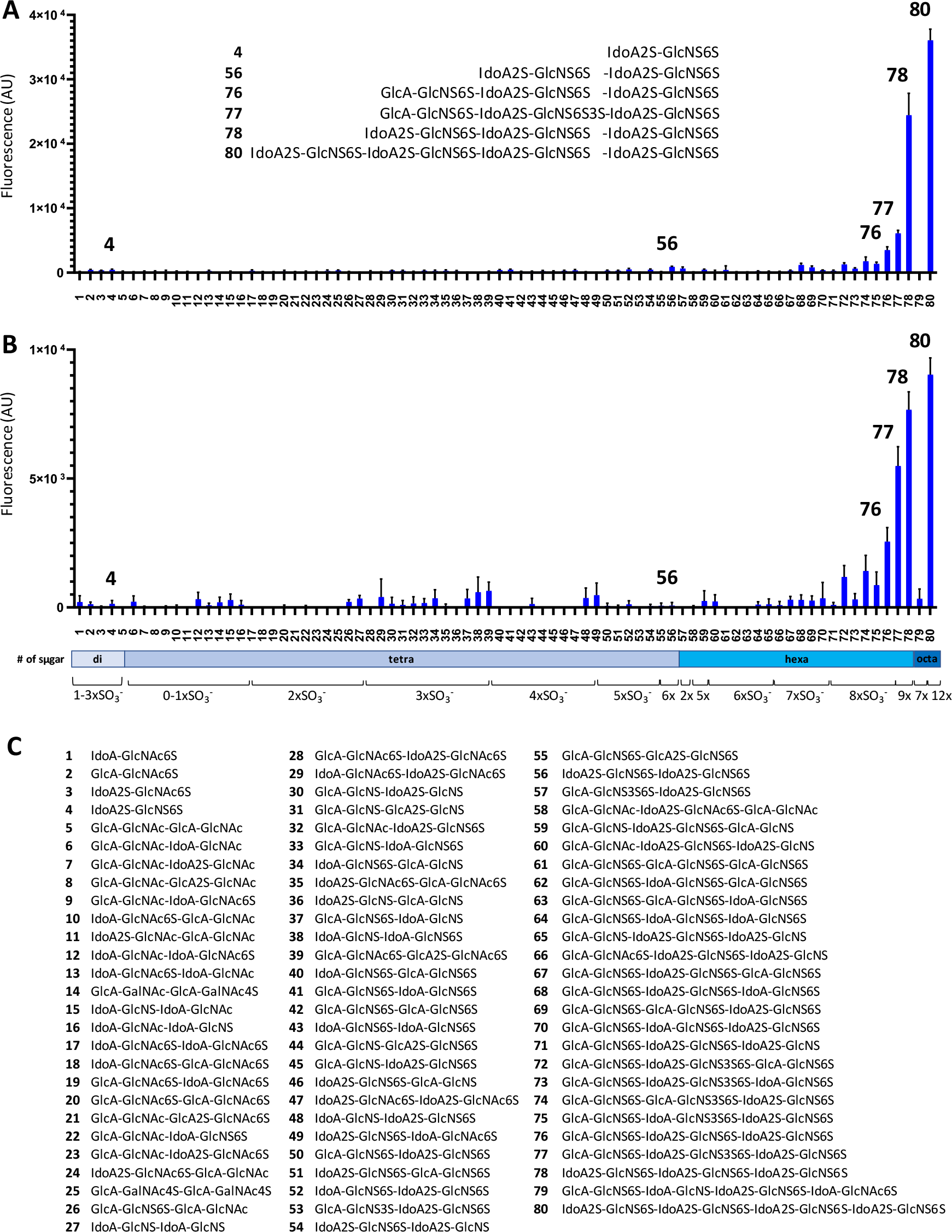
Binding of synthetic heparan sulfate oligosaccharides to SARS-CoV-2-spike and RBD by microarray. (A) Binding of SARS-CoV-2-spike (10 µg/mL) to the heparan sulfate microarray. The strongest binding structures are shown as inserts. (B) Binding of SARS-CoV2-RBD (30 µg/mL) on the heparan sulfate microarray. (C) Compounds numbering and structures of the heparan sulfate library.

Sub-arrays were incubated with different concentrations of SARS-CoV-2 RBD and spike protein in a binding buffer (pH 7.4, 20 mM Tris, 150 mM NaCl, 2 mM CaCl_2_, 2 mM MgCl_2_ with 1% BSA and 0.05% Tween-20) at room temperature for 1 h. After washing and drying, the subarrays were exposed to an anti-His antibody labeled with AlexaFluor® 647 for another hour, washed, dried and binding was detected by fluorescent scanning. To analyze the data, the compounds were arranged according to increasing backbone length, and within each group by increasing numbers of sulfates. Intriguingly, the proteins showed a strong preference for specific HS oligosaccharides (**Fig. 2A, B**). Furthermore, it was found that the RBD, monomeric spike protein, and trimeric spike protein exhibit similar binding patterns (**Fig. S1**). Compounds showing strong responsiveness (**76, 77, 78**, and **80**) are composed of tri-sulfated repeating units (IdoA2S-GlcNS6S). The binding is length-dependent and HS oligosaccharide **80** (IdoA2S-GlcNS6S)_4_ and **78** (IdoA2S-GlcNS6S)_3_ having four and three repeating units, respectively, showed the strongest binding. On the other hand, tetrasaccharide **56** (IdoA2S-GlcNS6S)_2_, which has the same repeating unit structure, gave very low responsiveness. A similar observation was made for disaccharide **4** (IdoA2S-GlcNS6S).

The structure-binding data shows that perturbations in the backbone or sulfation pattern led to substantial reductions in binding. The importance of the IdoA2S residue is highlighted by comparing hexasaccharides **78** with **76** in which a single IdoA2S in the distal disaccharide repeating unit is replaced with GlcA. This modification leads to a substantial reduction in responsiveness. Further replacements of IdoA2S with GlcA in compound **76** completely abolish binding, as evident for compounds **69, 67**, and **61**. The structure-activity data also showed that the 2-*O*-sulfates are crucial, and binding was lost when such functionalities were not present (**76** *vs*. **70, 68**, and **64**). Lack of one or more 6-*O*-sulfates also resulted in substantial reductions in binding (**76** *vs*. **71** and **65**). Although the SARS-CoV-2 spike and RBD showed similar selectivities, the binding of the spike appeared stronger and much higher fluorescent readings were observed at the same protein concentration.

Next, we examined whether HS oligosaccharide **80** can interfere in the interaction of the spike or RBD with immobilized heparin. Thus, the spike protein (150 nM) or RBD (2.4 µM) were pre-mixed with different concentrations of compound **80** and then used as analytes. The IC_50_ values were determined by non-linear fitting of Log(inhibitor) *vs*. response using variable slope (**Fig. S2**). The IC_50_ values for the spike protein and RBD are 38 nM and 264 nM, respectively.

To further determine the possible role of HS in the infection process, we examined the binding affinities of spike proteins to ACE2 and compared these with binding affinities for heparin. Biotinylated ACE2 was immobilized on a streptavidin-coated sensor chip and binding experiments were performed with different concentrations of the SARS-CoV-2 derived proteins. Representative sensorgrams for the RBD domain, monomeric spike protein, and trimeric spike protein are shown in **Fig. 3**. K_D_ values of 3.6 nM, 24.5 nM and 0.7 nM were determined using a 1:1 Langmuir binding model, respectively, which are in agreement with reported data.^28^ It shows convincingly that the RBD domain has a much higher affinity for ACE2 compared to that of heparin.

**Figure 3.**
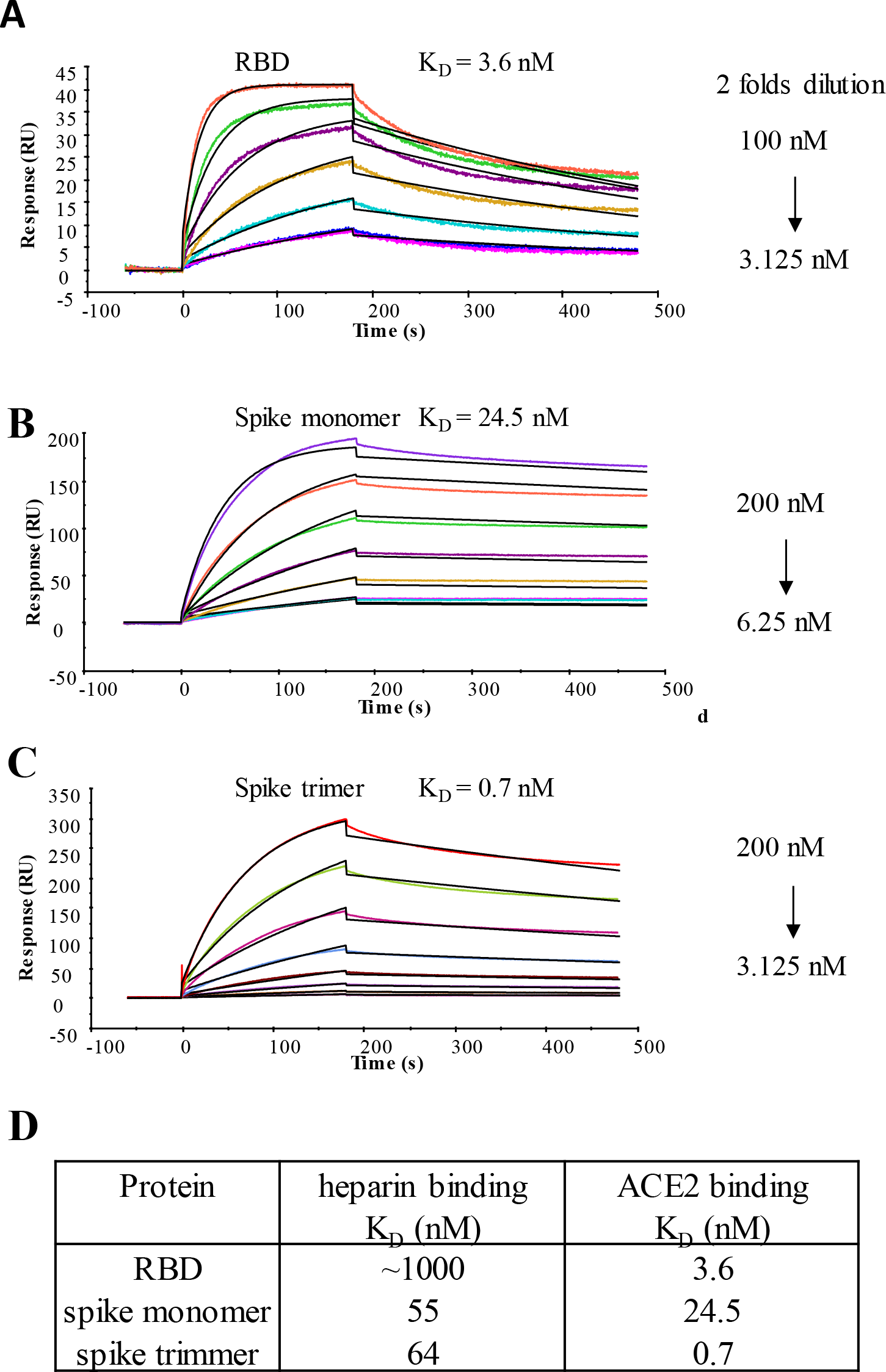
Sensorgrams representing the concentration-dependent kinetic analysis of the binding of immobilized ACE2 with SARS-CoV-2 derived proteins (A) RBD, (B) spike monomer, and (C) spike trimer. (D) Comparison of the K_D_ values of heparin binding and ACE2 binding to SARS-CoV-2 related proteins.

A number of reports have indicated that heparin and related compounds can block infection of cells by SARS-CoV-2. Therefore, we were compelled to investigate the molecular mechanisms by which heparin blocks viral entry.^2, 10, 13^ It is possible that the anti-viral properties of heparin are due to binding to the RBD domain thereby blocking the interaction with ACE2. Alternatively, heparin may interfere in the proteolytic processing of the spike protein thereby preventing membrane fusion. In this respect, the spike of SARS-CoV-2 contains a unique furin cleavage site, which is not present in other CoV’s, and has been proposed to contribute to high infectivity,^29^ because cleavage of the spike protein is a prerequisite for membrane fusion. Modeling studies have indicated that the furin cleavage site may harbor a binding site for HS.^27^ Finally, HS may function as an attachment factor and the addition of exogenous heparin may interfere in this process.

To examine whether heparin can interfere in binding of the spike to ACE2, we performed microarray experiments in which biotinylated Fc tagged ACE2 (50 μg/mL) was printed onto streptavidin coated microarray slides. The printing quality was confirmed by using a goat-anti-human Fc antibody conjugated with AlexaFluoro®647 (**Fig. S3A**). Next, His-tagged RBD and monomeric spike protein were premixed with different concentrations of heparin and binding of the proteins to immobilized ACE2 was accomplished by anti-His antibody. Soluble human ACE2 was used as positive control. Although, ACE2 efficiently inhibited RBD and spike binding (**Fig. S3 B, C**), no substantial changes in binding were observed in the presence of 10 μg/mL and 100 μg/mL of heparin (**Fig. 4 A, B**). Furthermore, we immobilized the RBD and monomeric spike proteins on ELISA plates and assayed the binding of ACE2 to the spike proteins in the presence or absence of heparin (**Fig. 4 C, D**). Soluble human ACE2 was used as a positive control, which as expected exhibited potent inhibition. At 100 μg/mL of heparin, no inhibition of binding was observed for either RBD or monomeric spike protein. These results indicate that heparin does not substantially interfere in the interaction of the spike with ACE2.

**Figure 4.**
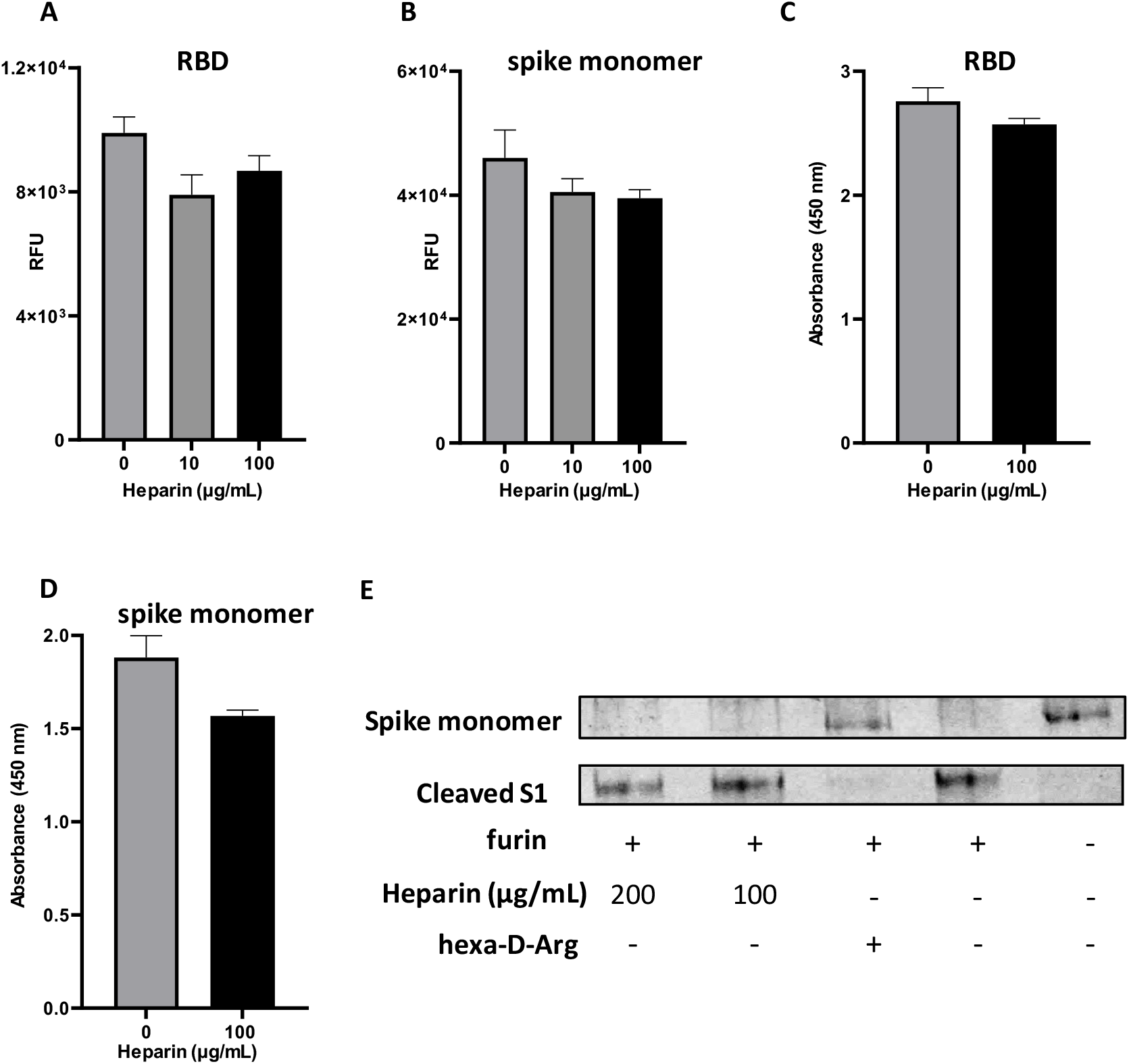
(A) Influence of heparin on the binding of His-tagged RBD or (B) His-tagged Spike monomer to biotinylated human ACE2 immobilized on streptavidin coated microarray slides. Detection of RBD and spike was accomplished using an anti-His antibody labeled with AlexaFluor 647. (C) Influence of heparin on the binding of biotinylated human ACE2 to RBD and (D) to immobilized spike monomer immobilized to high surface microtiter plates. Binding was detected by treatment with streptavidin-HRP followed by addition of a colorimetric HRP substrate. (E) Western Blot analysis of furin-mediated cleavage of spike monomer in the presence and absence of heparin or a known furin inhibitor (hexa-D-arginine).

To investigate whether the binding of heparin can hinder cleavage of the spike protein by furin, we exposed the monomeric spike protein to furin in the presence of different concentrations of heparin and examined protein cleavage by SDS-PAGE. The spike protein was readily cleaved by furin even in the presence of high concentration of heparin (400 μg/mL), while 50 μg/mL of a known furin inhibitor completely abolished cleavage.

It is also possible that heparin interferes in the initial attachment of the virus to the glycocalyx thereby preventing infection. Therefore, we examined the importance of HS for binding of trimeric RBD to relevant tissues.^30^ Ferrets are a susceptible animal model for SARS-CoV-2^31-32^ and closely related minks are easily infected on farms.^33^ Formalin-fixed, paraffin-embedded lung tissue slides resemble the complex membrane structures to which spike proteins need to bind before it can engage with ACE2 for cell entry. Expression of ACE2 was assessed using an ACE2 antibody allowing us to compare the binding with the SARS-CoV-RBD protein and binding localization and dependency on HS. The ACE2 antibody (**Fig. 5A**) and the RBD trimer bound efficiently to the ferret lung tissues (**Fig. 5B**). We also examined a commonly used heparan sulfate antibody, which bound efficiently to ferret lung tissue, indicating the omnipresence of HS. After overnight exposure to heparanase (HPSE), the ACE2 antibody staining was mostly unaffected, indicating HSPG-independent binding. On the other hand, the SARS-CoV-2 RBD trimer was not able to engage with the ferret lung tissue slide after HPSE treatment. No staining was observed with the heparin sulfate antibody (10E4), indicating all HS had been removed. Thus, these results indicate that HS is required for initial cell attachment before the spike can engage with ACE2.

**Figure 5.**
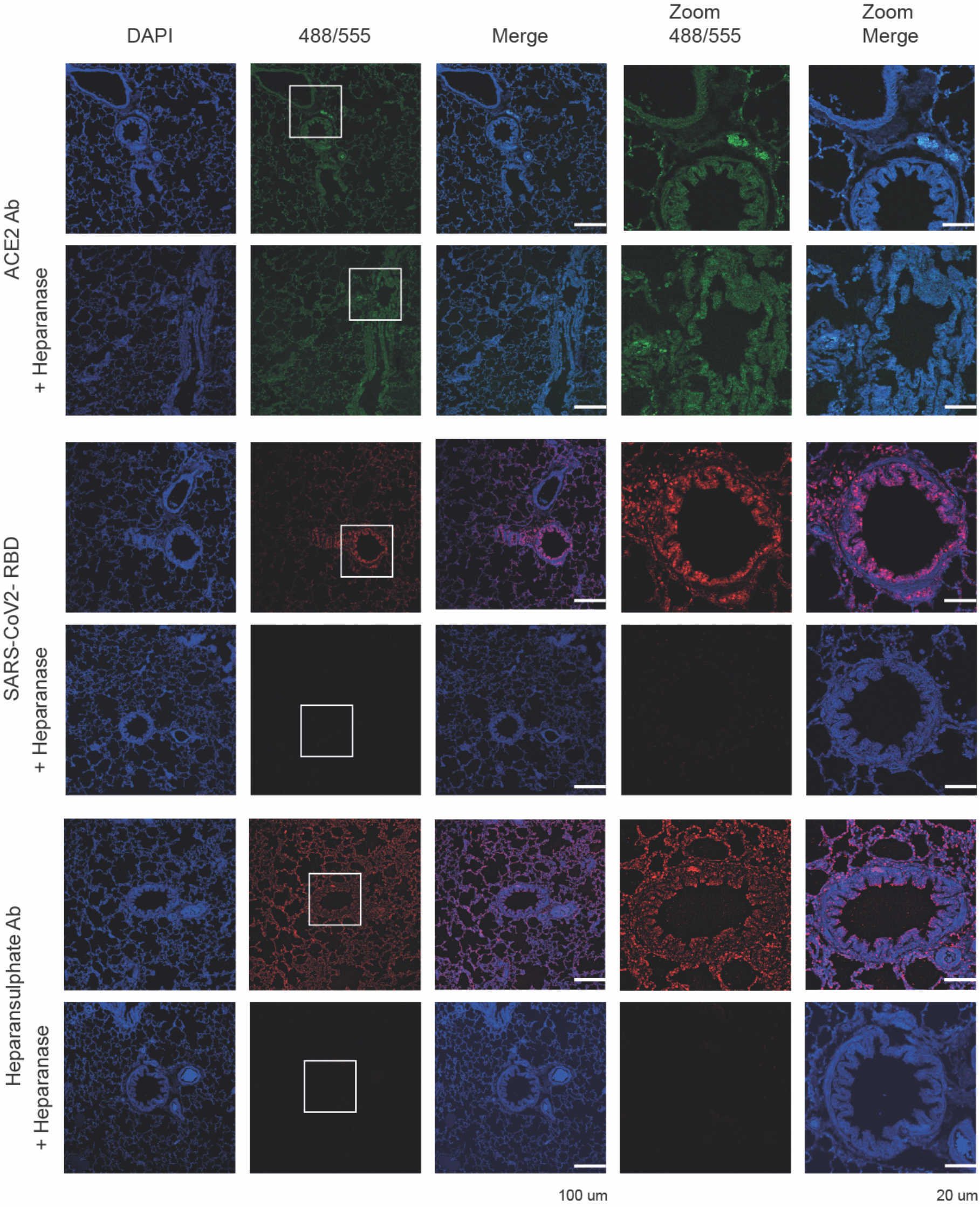
Binding of ACE2 antibody, SARS-CoV-2 RBD, and heparan sulfate antibody to ferret lung serial tissue slides. (A) ACE2 antibody staining without and after HPSE treatment. (B) SARS-CoV-2 RBD staining without and after HPSE treatment. (C) Heparan sulfate antibody (10E4) staining without and after HPSE treatment. HPSE treatment was achieved by overnight incubation of the tissues with HPSE (0.2 µg/mL) at 37 ^°^C.

## DISCUSSION AND CONCLUSIONS

The glycan microarray and SPR results indicate that the spike of SARS-CoV-2 can bind HS in a length- and sequence-dependent manner, and hexa- and octa-saccharides composed of IdoA2S-GlcNS6S repeating units have been defined as optimal ligands. The data supports a model in which the RBD of the spike confers sequence specificity and an additional HS binding site in the S1/S2 proteolytic cleavage site^9^ enhances the avidity of binding probably by non-specific interactions. In a BioRxiv preprint, we presented, for the first time, experimental support for such a model and subsequent papers have confirmed that the RBD harbors a HS binding site. Although IdoA2S-GlcNS6S sequons are abundantly present in heparin, it is a minor component of HS.^34^ Interestingly, it has been reported that the expression of the (GlcNS6S-IdoA2S)_3_ motif is highly regulated and plays a crucial role in cell behavior and disease including endothelial cell activation.^35^ Severe thrombosis in COVID-19 patients is associated with endothelial dysfunction^36^ and a connection may exist between SARS-CoV-2’s ability to bind to HS and thrombotic disorder. It is also possible that HS is a determinant of the cell- and tissue tropism.

A number of reports have shown that heparin and related products can block infection by pseudotyped virus or authentic SARS-CoV-2 virus.^12-14, 27^ We explored the possibility that binding of heparin blocks the RBD from interacting with ACE2. However, in two experimental formats such properties were not observed. We found that the affinity of the RBD for heparin is much lower than that for ACE2, providing a rationale for the inability of heparin to inhibit the binding between RBD or spike with ACE2. One computational study has indicated that ACE2 and HS bind to the same region of the RBD.^27^ Another docking study located the HS binding site adjacent to the ACE2-binding site and inferred a model in which a ternary complex is formed between RBD, HS and ACE2.^14^ Further studies are required to determine the exact location of the HS binding site, which in turn may provide a better understanding of the interplay between binding of spike with ACE2 and heparin.

We employed physiological relevant tissues to explore the importance of HS for SARS-CoV-2 adhesion and demonstrated that HPSE treatment greatly reduces RBD binding but not that of ACE2. The data supports a model in which HS functions as a host attachment factor that facilitates SARS-CoV-2 infection.

The current clinical guidelines call for the use of unfractionated heparin or low molecular weight heparin (LMWH) for the treatment of all COVID-19 patients for systemic clotting in the absences of contradictions.^37-38^ Heparin treatment may have additional benefits and may compete with the binding of the spike protein to cell surface HS thereby preventing infectivity. Our data suggest that non-coagulating heparin or HS preparations can be developed that reduce cell binding and infectivity without a risk of causing bleeding. In this respect, administration of heparin requires great care because its anticoagulant activity can result in excessive bleeding. Antithrombin III (AT-III), which confers anticoagulant activity, binds a specific pentasaccharide GlcNAc(6S)-GlcA-GlcNS(3S)(6S)-IdoA2S-GlcNS(6S) embedded in HS or heparin. Removal of the sulfate at C-3 of *N*-sulfoglucosamine (GlcNS3S) of the pentasaccharide results in a 10^5^-fold reduction in binding affinity.^39^ Importantly, such a functionality is not present in the identified HS ligand of SARS-CoV-2 spike, and therefore compounds can be developed that can inhibit cell binding, but do not interact with ATIII. As a result, such preparations can be used at higher doses without causing adverse side effects. Our data also shows that multivalent interactions of the spike with HS results in high avidity of binding. This observation provides opportunities to develop glycopolymers modified by HS oligosaccharides as inhibitors of SARS-CoV-2 cell binding to prevent or treat COVID-19.

## ACKNOWLEDGMENTS

This research was supported by the National Institutes of Health (P41GM103390 and R01HL151617 to G.-J.B.). R.P.dV is a recipient of an ERC Starting Grant from the European Commission (802780) and a Beijerinck Premium of the Royal Dutch Academy of Sciences. We thank Sander Herfst (Department of Viroscience, Erasmus Medical Center) for the ferret tissues and Gavin Wright (Addgene) for providing HPSE-bio-His (Plasmid #53407). Plasmids for expression of SARS-CoV-2 spike and RBD proteins were provided by Dr. Florian Krammer (Icahn School of Medicine at Mount Sinai, produced under NIAID CEIRS contract HHSN272201400008C). Production of recombinant proteins was supported by NIAID Centers of Excellence for Influenza Research and Surveillance (CEIRS) contract HHSN272201400004C to S.M.T.

## REFERENCES

1. Dimitrov, D. S., Virus entry: molecular mechanisms and biomedical applications. Nat. Rev. Microbiol. 2004, 2 (2), 109–122.

2. Walls, A. C.; Park, Y.-J.; Tortorici, M. A.; Wall, A.; McGuire, A. T.; Veesler, D., Structure, function, and antigenicity of the SARS-CoV-2 spike glycoprotein. Cell 2020, 181 (2), 281-292.e6.

3. Li, F.; Li, W.; Farzan, M.; Harrison, S. C., Structure of SARS coronavirus spike receptor-binding domain complexed with receptor. Science 2005, 309 (5742), 1864–1868.

4. Monteil, V.; Kwon, H.; Prado, P.; Hagelkrüys, A.; Wimmer, R. A.; Stahl, M.; Leopoldi, A.; Garreta, E.; Hurtado del Pozo, C.; Prosper, F.; Romero, J. P.; Wirnsberger, G.; Zhang, H.; Slutsky, A. S.; Conder, R.; Montserrat, N.; Mirazimi, A.; Penninger, J. M., Inhibition of SARS-CoV-2 infections in engineered human tissues using clinical-grade soluble human ACE2. Cell 2020, 181 (1), 1–9.

5. Li, W.; Hulswit, R. J. G.; Widjaja, I.; Raj, V. S.; McBride, R.; Peng, W.; Widagdo, W.; Tortorici, M. A.; van Dieren, B.; Lang, Y.; van Lent, J. W. M.; Paulson, J. C.; de Haan, C. A. M.; de Groot, R. J.; van Kuppeveld, F. J. M.; Haagmans, B. L.; Bosch, B.-J., Identification of sialic acid-binding function for the Middle East respiratory syndrome coronavirus spike glycoprotein. Proc. Natl. Acad. Sci. 2017, 114 (40), E8508–E8517.

6. Milewska, A.; Zarebski, M.; Nowak, P.; Stozek, K.; Potempa, J.; Pyrc, K., Human coronavirus NL63 utilizes heparan sulfate proteoglycans for attachment to target cells. J. Virol. 2014, 88 (22), 13221–13230.

7. Lang, J.; Yang, N.; Deng, J.; Liu, K.; Yang, P.; Zhang, G.; Jiang, C., Inhibition of SARS pseudovirus cell entry by lactoferrin binding to heparan sulfate proteoglycans. PLoS One 2011, 6 (8), e23710.

8. Mycroft-West, C.; Su, D.; Elli, S.; Li, Y.; Guimond, S.; Miller, G.; Turnbull, J.; Yates, E.; Guerrini, M.; Fernig, D.; Lima, M.; Skidmore, M., The 2019 coronavirus (SARS-CoV-2) surface protein (Spike) S1 receptor binding domain undergoes conformational change upon heparin binding. bioRxiv 2020, 2020.02.29.971093.

9. Kim, S. Y.; Jin, W.; Sood, A.; Montgomery, D. W.; Grant, O. C.; Fuster, M. M.; Fu, L.; Dordick, J. S.; Woods, R. J.; Zhang, F.; Linhardt, R. J., Glycosaminoglycan binding motif at S1/S2 proteolytic cleavage site on spike glycoprotein may facilitate novel coronavirus (SARS-CoV-2) host cell entry. bioRxiv 2020, 2020.04.14.041459.

10. Tang, T.; Bidon, M.; Jaimes, J. A.; Whittaker, G. R.; Daniel, S., Coronavirus membrane fusion mechanism offers a potential target for antiviral development. Antiviral Res. 2020, 178, 104792.

11. Partridge, L. J.; Urwin, L.; Nicklin, M. J. H.; James, D. C.; Green, L. R.; Monk, P. N., ACE2-independent interaction of SARS-CoV-2 spike protein to human epithelial cells can be inhibited by unfractionated heparin. bioRxiv 2020, 2020.05.21.107870.

12. Guimond, S. E.; Mycroft-West, C. J.; Gandhi, N. S.; Tree, J. A.; Buttigieg, K. R.; Coombes, N.; Nystrom, K.; Said, J.; Setoh, Y. X.; Amarilla, A.; Modhiran, N.; Julian Sng, D. J.; Chhabra, M.; Watterson, D.; Young, P. R.; Khromykh, A. A.; Lima, M. A.; Fernig, D. G.; Su, D.; Yates, E. A.; Hammond, E.; Dredge, K.; Carroll, M. W.; Trybala, E.; Bergstrom, T.; Ferro, V.; Skidmore, M. A.; Turnbull, J. E., Pixatimod (PG545), a clinical-stage heparan sulfate mimetic, is a potent inhibitor of the SARS-CoV-2 virus. bioRxiv 2020, 2020.06.24.169334.

13. Mycroft-West, C. J.; Su, D.; Pagani, I.; Rudd, T. R.; Elli, S.; Guimond, S. E.; Miller, G.; Meneghetti, M. C. Z.; Nader, H. B.; Li, Y.; Nunes, Q. M.; Procter, P.; Mancini, N.; Clementi, M.; Bisio, A.; Forsyth, N. R.; Turnbull, J. E.; Guerrini, M.; Fernig, D. G.; Vicenzi, E.; Yates, E. A.; Lima, M. A.; Skidmore, M. A., Heparin inhibits cellular invasion by SARS-CoV-2: structural dependence of the interaction of the surface protein (spike) S1 receptor binding domain with heparin. bioRxiv 2020, 2020.04.28.066761.

14. Clausen, T. M.; Sandoval, D. R.; Spliid, C. B.; Pihl, J.; Perrett, H. R.; Painter, C. D.; Narayanan, A.; Majowicz, S. A.; Kwong, E. M.; McVicar, R. N.; Thacker, B. E.; Glass, C. A.; Yang, Z.; Torres, J. L.; Golden, G. J.; Bartels, P. L.; Porell, R. N.; Garretson, A. F.; Laubach, L.; Feldman, J.; Yin, X.; Pu, Y.; Hauser, B. M.; Caradonna, T. M.; Kellman, B. P.; Martino, C.; Gordts, P. L. S. M.; Chanda, S. K.; Schmidt, A. G.; Godula, K.; Leibel, S. L.; Jose, J.; Corbett, K. D.; Ward, A. B.; Carlin, A. F.; Esko, J. D., SARS-CoV-2 Infection Depends on Cellular Heparan Sulfate and ACE2. Cell 2020, 183 (4), 1043-1057.e15.

15. Bishop, J. R.; Schuksz, M.; Esko, J. D., Heparan sulphate proteoglycans fine-tune mammalian physiology. Nature 2007, 446 (7139), 1030–1037.

16. Cagno, V.; Tseligka, E. D.; Jones, S. T.; Tapparel, C., Heparan sulfate proteoglycans and viral attachment: true receptors or adaptation bias? Viruses 2019, 11 (7), 596.

17. de Haan, C. A. M.; Haijema, B. J.; Schellen, P.; Wichgers Schreur, P.; te Lintelo, E.; Vennema, H.; Rottier, P. J. M., Cleavage of group 1 coronavirus spike proteins: how furin cleavage is traded off against heparan sulfate binding upon cell culture adaptation. J. Virol. 2008, 82 (12), 6078–6083.

18. de Haan, C. A. M.; Li, Z.; te Lintelo, E.; Bosch, B. J.; Haijema, B. J.; Rottier, P. J. M., Murine coronavirus with an extended host range uses heparan sulfate as an entry receptor. J. Virol. 2005, 79 (22), 14451–14456.

19. Sarrazin, S.; Lamanna, W. C.; Esko, J. D., Heparan sulfate proteoglycans. Cold Spring Harb. Perspect. Biol. 2011, 3 (7), a004952.

20. Xu, D.; Esko, J. D., Demystifying heparan sulfate–protein interactions. Annu. Rev. Biochem 2014, 83 (1), 129–157.

21. Kamhi, E.; Joo, E. J.; Dordick, J. S.; Linhardt, R. J., Glycosaminoglycans in infectious disease. Biol. Rev. 2013, 88 (4), 928–943.

22. García, B.; Merayo-Lloves, J.; Martin, C.; Alcalde, I.; Quirós, L. M.; Vazquez, F., Surface proteoglycans as mediators in bacterial pathogens infections. Front. Microbiol. 2016, 7, 220.

23. Zong, C.; Venot, A.; Li, X.; Lu, W.; Xiao, W.; Wilkes, J.-S. L.; Salanga, C. L.; Handel, T. M.; Wang, L.; Wolfert, M. A.; Boons, G.-J., Heparan sulfate microarray reveals that heparan sulfate–protein binding exhibits different ligand requirements. J. Am. Chem. Soc. 2017, 139 (28), 9534–9543.

24. Arungundram, S.; Al-Mafraji, K.; Asong, J.; Leach, F. E.; Amster, I. J.; Venot, A.; Turnbull, J. E.; Boons, G.-J., Modular Synthesis of Heparan Sulfate Oligosaccharides for Structure−Activity Relationship Studies. J. Am. Chem. Soc. 2009, 131 (47), 17394–17405.

25. Stadlbauer, D.; Amanat, F.; Chromikova, V.; Jiang, K.; Strohmeier, S.; Arunkumar, G. A.; Tan, J.; Bhavsar, D.; Capuano, C.; Kirkpatrick, E.; Meade, P.; Brito, R. N.; Teo, C.; McMahon, M.; Simon, V.; Krammer, F. SARS-CoV-2 seroconversion in humans: A detailed protocol for a serological assay, antigen production, and test setup. Curr. Protoc. Microbiol. 2020, 57 (1), e100..

26. Amanat, F.; Stadlbauer, D.; Strohmeier, S.; Nguyen, T. H. O.; Chromikova, V.; McMahon, M.; Jiang, K.; Arunkumar, G. A.; Jurczyszak, D.; Polanco, J.; Bermudez-Gonzalez, M.; Kleiner, G.; Aydillo, T.; Miorin, L.; Fierer, D. S.; Lugo, L. A.; Kojic, E. M.; Stoever, J.; Liu, S. T. H.; Cunningham-Rundles, C.; Felgner, P. L.; Moran, T.; Garcia-Sastre, A.; Caplivski, D.; Cheng, A. C.; Kedzierska, K.; Vapalahti, O.; Hepojoki, J. M.; Simon, V.; Krammer, F. A serological assay to detect SARS-CoV-2 seroconversion in humans. Nat. Med. 2020, 26 (7), 1033–1036.

27. Kim, S. Y.; Jin, W.; Sood, A.; Montgomery, D. W.; Grant, O. C.; Fuster, M. M.; Fu, L.; Dordick, J. S.; Woods, R. J.; Zhang, F.; Linhardt, R. J., Characterization of heparin and severe acute respiratory syndrome-related coronavirus 2 (SARS-CoV-2) spike glycoprotein binding interactions. Antiviral Res. 2020, 181, 104873.

28. Shang, J.; Ye, G.; Shi, K.; Wan, Y.; Luo, C.; Aihara, H.; Geng, Q.; Auerbach, A.; Li, F., Structural basis of receptor recognition by SARS-CoV-2. Nature 2020, 581 (7807), 221–224.

29. Xia, S.; Lan, Q.; Su, S.; Wang, X.; Xu, W.; Liu, Z.; Zhu, Y.; Wang, Q.; Lu, L.; Jiang, S., The role of furin cleavage site in SARS-CoV-2 spike protein-mediated membrane fusion in the presence or absence of trypsin. Signal. Transduct. Target. Ther. 2020, 5 (1), 92.

30. Bouwman, K. M.; Tomris, I.; Turner, H. L.; van der Woude, R.; Bosman, G. P.; Rockx, B.; Herfst, S.; Haagmans, B. L.; Ward, A. B.; Boons, G.-J.; de Vries, R. P., Multimerization-and glycosylation-dependent receptor binding of SARS-CoV-2 spike proteins. bioRxiv 2020, 2020.09.04.282558.

31. Kim, Y.-I.; Kim, S.-G.; Kim, S.-M.; Kim, E.-H.; Park, S.-J.; Yu, K.-M.; Chang, J.-H.; Kim, E. J.; Lee, S.; Casel, M. A. B.; Um, J.; Song, M.-S.; Jeong, H. W.; Lai, V. D.; Kim, Y.; Chin, B. S.; Park, J.-S.; Chung, K.-H.; Foo, S.-S.; Poo, H.; Mo, I.-P.; Lee, O.-J.; Webby, R. J.; Jung, J. U.; Choi, Y. K., Infection and rapid transmission of SARS-CoV-2 in ferrets. Cell Host Microbe 2020, 27 (5), 704-709.e2.

32. Richard, M.; Kok, A.; de Meulder, D.; Bestebroer, T. M.; Lamers, M. M.; Okba, N. M. A.; Fentener van Vlissingen, M.; Rockx, B.; Haagmans, B. L.; Koopmans, M. P. G.; Fouchier, R. A. M.; Herfst, S., SARS-CoV-2 is transmitted via contact and via the air between ferrets. Nat. Comm. 2020, 11 (1), 3496.

33. Oreshkova, N.; Molenaar, R. J.; Vreman, S.; Harders, F.; Oude Munnink, B. B.; Hakze-van der Honing, R. W.; Gerhards, N.; Tolsma, P.; Bouwstra, R.; Sikkema, R. S.; Tacken, M. G.; de Rooij, M. M.; Weesendorp, E.; Engelsma, M. Y.; Bruschke, C. J.; Smit, L. A.; Koopmans, M.; van der Poel, W. H.; Stegeman, A., SARS-CoV-2 infection in farmed minks, the Netherlands, April and May 2020. Eurosurveillance 2020, 25 (23), 2001005.

34. Rabenstein, D. L., Heparin and heparan sulfate: structure and function. Nat. Prod. Rep. 2002, 19 (3), 312–331.

35. Smits, N. C.; Kurup, S.; Rops, A. L.; ten Dam, G. B.; Massuger, L. F.; Hafmans, T.; Turnbull, J. E.; Spillmann, D.; Li, J.-p.; Kennel, S. J.; Wall, J. S.; Shworak, N. W.; Dekhuijzen, P. N. R.; van der Vlag, J.; van Kuppevelt, T. H., The heparan sulfate motif (GlcNS6S-IdoA2S)3, common in heparin, has a strict topography and is involved in cell behavior and disease. J. Biol. Chem. 2010, 285 (52), 41143–41151.

36. Sardu, C. G.J.; Morelli, M.B.; Wang, X.; Marfella, R.; Santulli, G., Is COVID-19 an endothelial disease? Clinical and basic evidence. Preprints 2020, 2020040204.

37. Tang, N.; Bai, H.; Chen, X.; Gong, J.; Li, D.; Sun, Z., Anticoagulant treatment is associated with decreased mortality in severe coronavirus disease 2019 patients with coagulopathy. J. Thromb. Haemost. 2020, 18 (5), 1094–1099.

38. Thachil, J.; Tang, N.; Gando, S.; Falanga, A.; Cattaneo, M.; Levi, M.; Clark, C.; Iba, T., ISTH interim guidance on recognition and management of coagulopathy in COVID-19. J. Thromb. Haemost. 2020, 18 (5), 1023–1026.

39. Thacker, B. E.; Xu, D.; Lawrence, R.; Esko, J. D., Heparan sulfate 3-O-sulfation: a rare modification in search of a function. Matrix Biol. 2014, 35, 60–72.

