## Supporting Information for "Heparan sulfate proteoglycans as attachment factor for SARS-CoV-2"

### Table of Contents

**Materials** 2

**Methods** 2

Expression of SARS-CoV-2-RBD 2

Microarray printing and screening 2

Preparation of biotinylated heparin 3

Surface plasma resonances experiments 3

Competition experiments using immobilized ACE2 on streptavidin coated slides 3

The ability of heparin to inhibit spike protein RBD or spike protein monomer by ELISA 4

Furin cleavage assay 4

Tissue binding experiments 4

**Supplementary Figures** 6

Figure S1 6

Figure S2 8

Figure S3 9

**References** 10

### Materials

The SARS-CoV-2-spike (S1+S2) was obtained from Sino Biological (40589-V08B1). The Alexa Fluor 647 anti-His tag antibody was obtained from BioLegend (#652513). High grade heparin (HG-Hep, Mw 15,700) for SPR was obtained from Iduron Ltd, UK (#HEP-HG 100). Furin was obtained from New England Biolabs (P8077). Furin inhibitor II (hexa-D-Arginine) was obtained from Sigma-Aldrich (SCP0148). The SARS-CoV-2 (COVID-19) inhibitor screening kit (EP-105) and the SARS-CoV-2 inhibitor (Human ACE2; AC2-NA005) were from ACRO Biosystems.

### Methods

#### Expression of SARS-CoV-2 RBD and Spike proteins

The RBD modified by an His6-tag and trimerized, soluble spike with His6-tag were expressed in HEK cells and purified as previously described.[1,2] The RBD trimer used in the tissue binding experiments was expressed as previously described.[3]

#### Microarray printing and screening

Aminopentyl modified HS oligosaccharides were printed on NHS-ester activated glass slides (NEXTERION® Slide H, Schott Inc.) using a Scienion sciFLEXARRAYER S3 non-contact microarray equipped with a Scienion PDC80 nozzle (Scienion Inc.). Individual samples were dissolved in sodium phosphate buffer (50 µL, 0.225 M, pH 8.5) at a concentration of 100 µM and were printed in replicates of 10 with a spot volume of ~ 400 pL at 20 °C and 50% humidity. Each slide has 24 subarrays in a 3x8 layout. After printing, slides were incubated in a humidity chamber for 8 h and then blocked for 30 min in a Tris buffer (pH 9.0, 50 mM) containing 5 mM ethanolamine at 40 °C. Blocked slides were rinsed with DI water, spun dry, and kept in a desiccator at room temperature for future use.

Screening was performed by incubating the slides with a protein solution at the indicated time, followed by washing and drying. A typical washing procedure includes sequentially dipping the glass slide in TSM wash buffer (2 min, containing 0.05 % Tween 20), TSM buffer (2 min) and, water (2 x 2 min), followed by centrifugation. For SARS-CoV-2-RBD and SARS-CoV-2-spike protein screening, the slides were incubated with SARS-CoV-2-RBD (10, 30, and 100 µg/mL in TSMBB) or SARS-CoV-2-spike (3, 10, and 30 µg/mL in TSMBB) for 1 h, followed by washing and incubation with a solution of AlexaFluor® 647 conjugated anti-His antibody (10 µg/mL). After washing and drying, the slides were scanned using a GenePix 4000B microarray scanner (Molecular Devices) at the appropriate excitation wavelength with a resolution of 5 µM. Optimum gains and PMT values were employed for the scanning ensuring that all signals were within the linear range of the scanner's detector and there was no saturation of signals. The images were analyzed using GenePix Pro 7 software (version 7.2.29.2, Molecular Devices). The data was analyzed with a home written Excel macro. The highest and the lowest value of the total fluorescence intensity of the replicate spots were removed, and the remaining values were used to provide the mean value and standard deviation.

**Preparation of biotinylated heparin**

High grade heparin (HG-Hep, Mw 15,700, Iduron, UK) was biotinylated following a literature procedure.[4] Briefly, HG-Hep (0.5 mg), biotin-$(PEG)_3$-$NH_2$ (0.5 mg, Thermo Scientific, USA) and sodium cyanoborohydride (2.5 mg, $NaBH_3CN$, AminoLink™ reductant, Thermo Scientific, USA) were combined, dissolved in water (100 µL) and stirred at 70 °C for 24 h. Another portion of $NaBH_3CN$ (2.5 mg) was added and stirring continued for additional 24 h. To remove excess of biotin linker and reductant, the reaction mixture was centrifuged over a spin filter (Mw cutoff 3000) and exchanged with water (2x1 mL, Milli-Q®), concentrated (200 µL) and lyophilized to obtain the desired Biotin-HG-Hep as white fluffy powder. The product was aliquoted and stored at -80 °C.

**Surface plasma resonances experiments**

A streptavidin coated sensor chip was prepared from a CM5 chip by standard amine coupling using an amine coupling kit (Biacore Inc. - GE Healthcare). Briefly, the surface was activated using freshly mixed *N*-hydroxysuccinimide (NHS; 100 mM) and 1-(3-dimethylaminopropyl)-ethylcarbodiimide (EDC; 391 mM) (1/1, v/v) in water. The remaining active esters were quenched by aqueous ethanolamine (1.0 M; pH 8.5). Next, streptavidin (50 µg/mL) in aqueous NaOAc (10 mM, pH 4.5) was passed over the chip surface until a ligand density of approximately 3,000 RU was achieved. Next, biotin-HG-Hep (1 µM) was passed over one of the flow channels at a flow rate of 5 µL/min for 2 min resulting in a response of 56 RU. Next, the reference and modified flow cells were washed with three consecutive injections of 1 min with 1 M NaCl in 50 mM NaOH. HBS-EP (0.01 M HEPES, 150 mM NaCl, 3 mM EDTA, 0.005% polysorbate 20; pH 7.4) was used as the running buffer for the immobilization, kinetic studies of the interaction of HG-Hep with SARS-CoV-2-RBD and spike protein (S1+S2), and inhibition studies of immobilized HG-Hep with synthetic GAGs oligosaccharides. Analytes were dissolved in running buffer and a flow rate of 30 µL/min was employed for association and dissociation at a constant temperature of 25 ºC. A 60 s injection of 0.25% sodium dodecyl sulfonate (SDS) at a flow rate of 30 µL/min was used for regeneration and to achieve prior baseline status. Using Biacore T100 evaluation software, the response curves of various analyte concentrations were globally fitted to the 1:1 binding model.

Human angiotensin-converting enzyme 2 (ACE2) with Fc tag (Sino biology 10108-H02H) (5 µg/mL in HBS-P buffer) was captured to a human IgG Fc antibody chip at a flow rate of 5 µL/min for 300 s before the injection of analytes. SARSCov2RBD and spike protein were diluted in running buffer and a flow rate of 30 µL/min was employed for association and dissociation at a constant temperature of 25 ºC. A 30 s injection of 10 mM glycine (pH 1.5) at a flow rate of 50 µL/min was used for regeneration and achieved prior baseline status. Using Biacore T100 evaluation software, the response curves of various analyte concentrations were globally fitted to the 1:1 binding model.

**Competition experiments using immobilized ACE2 on streptavidin coated slides**

Biotinylated ACE2 (ACRO Biosystems, AC2-H82F9) was printed on streptavidin coated glass slides (SuperStreptavidin microarray substrate slides, ArrayIt Inc) at 12.5, 25, and 50 µg/mL using a Scienion sciFLEXARRAYER S3 non-contact microarray equipped with a Scienion PDC80 nozzle (Scienion Inc) in replicates of six. The average drop volume was 400 pL. The slides were stored at 4 °C after printing and were blocked with TSM binding buffer (20 mM Tris·Cl, pH 7.4, 150 mM NaCl, 2 mM CaCl2, and 2 mM MgCl2, 0.05% Tween-20, 1% BSA) for 1 h at 4 °C prior to use.

The protein (SARS-CoV-2 RBD-His or spike monomer-His 2.5 µg/mL) was premixed with AlexaFluor 647 conjugated anti-His antibody (2.5 µg/mL), followed by addition of human ACE2 as SARS-CoV-2 inhibitor (ACRO Biosystems, AC2-NA005) at various concentrations. The mixture was incubated at 4°C for 15 min before added to the microarray surface. The microarray slide was incubated at room temperature for 45 min, washed with TSM washing buffer, TSM buffer, water, and spun dried. The washing steps was carried out with the cassette attached to protect the unused blocks. After staining, the slides were scanned using a GenePix 4000B microarray scanner (Molecular Devices) at the appropriate excitation wavelength with a resolution of 10 µM. Optimum gains and PMT values were employed in the scanning to ensure that all the signals were within the liner range of the scanner's detector and there was no saturation of signals. The image was analyzed using GenePix Pro 7 software (version 7.2.29.2, Molecular Devices). The data was analyzed using a home written Excel macro. The highest and the lowest value of the total fluorescence intensity of the six replicates spots were removed, and the remaining four values were used to provide the mean value and standard deviation.

**The ability of heparin to inhibit spike protein RBD or spike protein monomer by ELISA**

ACE2 binding was determined by a colorimetric ELISA assay (SARS-CoV-2 inhibitor screening kit, ACROBiosytems). The assay was performed according to the manufacturer's instruction. Briefly, desired spike protein component was immobilized (2.5 µg/mL, 100 µL/well) at a high binding surface microplate (clear bottom) at 4 °C for 16 h. The microwells were then incubated with biotinylated human ACE2 (1 µg/mL, 50 µL/well) either in the presence of heparin (HG-heparin, 200 µg/mL, 50 µL/well) or in the absence of heparin (dilution buffer, 50 µL/well) at 37 °C for 1 h. Next, streptavidin-HRP (0.1 µg/mL, 100 µL/well) was added to each microwell and incubated at 37 °C for 1 h, followed by treatment with colorimetric HRP substrate (TMB One Component, SouthernBiotech, 200 µL/well) at 37 °C for 20 min. The reaction was terminated by the addition of a stop solution (1 M sulfuric acid, 50 µL/well) and absorbance was measured at 450 nm (POLARstar Optima, BMG labtech). The absorbance values were corrected for measured blank value and plotted using Prism 9 software (GraphPad Software, Inc.), bars represent the mean ± SD for each treatment. The assay was performed (in triplicate) at least two times.

**Furin cleavage assay**

Furin cleavage reactions were performed with 2 µg of SARS-CoV-2 spike protein, 1 µL furin and different concentrations of heparin or furin inhibitor II in 25 mM Tris-HCl buffer (pH 7.0) with 2 mM of $CaCl_2$ in a total volume of 10 µL. The reactions were incubated at 37 °C for 1 h before the reaction was loaded on an SDS-gel for analysis.

**Tissue binding experiments**

Serial sections of formalin-fixed, paraffin-embedded ferret lungs were obtained from the Department of Veterinary Pathobiology, Faculty of Veterinary Medicine, Utrecht

University and the Department of Viroscience, Erasmus University, The Netherlands, respectively. Tissue sections were rehydrated in a series of alcohol from 100%, 96% to 70%, and lastly in distilled water. Tissues slides were boiled in citrate buffer pH 6.0 for 10 min at 900 kW in a microwave for antigen retrieval and washed in PBS-T three times. Endogenous peroxidase activity was blocked with 1% hydrogen peroxide for 30 min. Tissues were subsequently incubated with 3% BSA in PBS-T overnight at 4 °C. The next day, the purified viral spike proteins (50 µg/mL), a rabbit antibody against ACE2 (5 µg/mL; Abcam 272690) and a mouse antibody against heparan sulphate (10 µg/mL; Abcam 370255-S) were added to the tissues for 1 h at RT. With rigorous washing steps in between, the proteins were detected using goat anti mouse (AlexaFluor 555) and anti-rabbit (AlexaFluor 488) antibodies at 400 ng/mL (Life Biosciences). Where indicated, tissues were incubated overnight at 37 °C with 0.2 µg/mL heparanase (HPSE). HPSE was expressed in HEK 293T cells using Addgene plasmid #53407.[5]

### Supplementary Figures

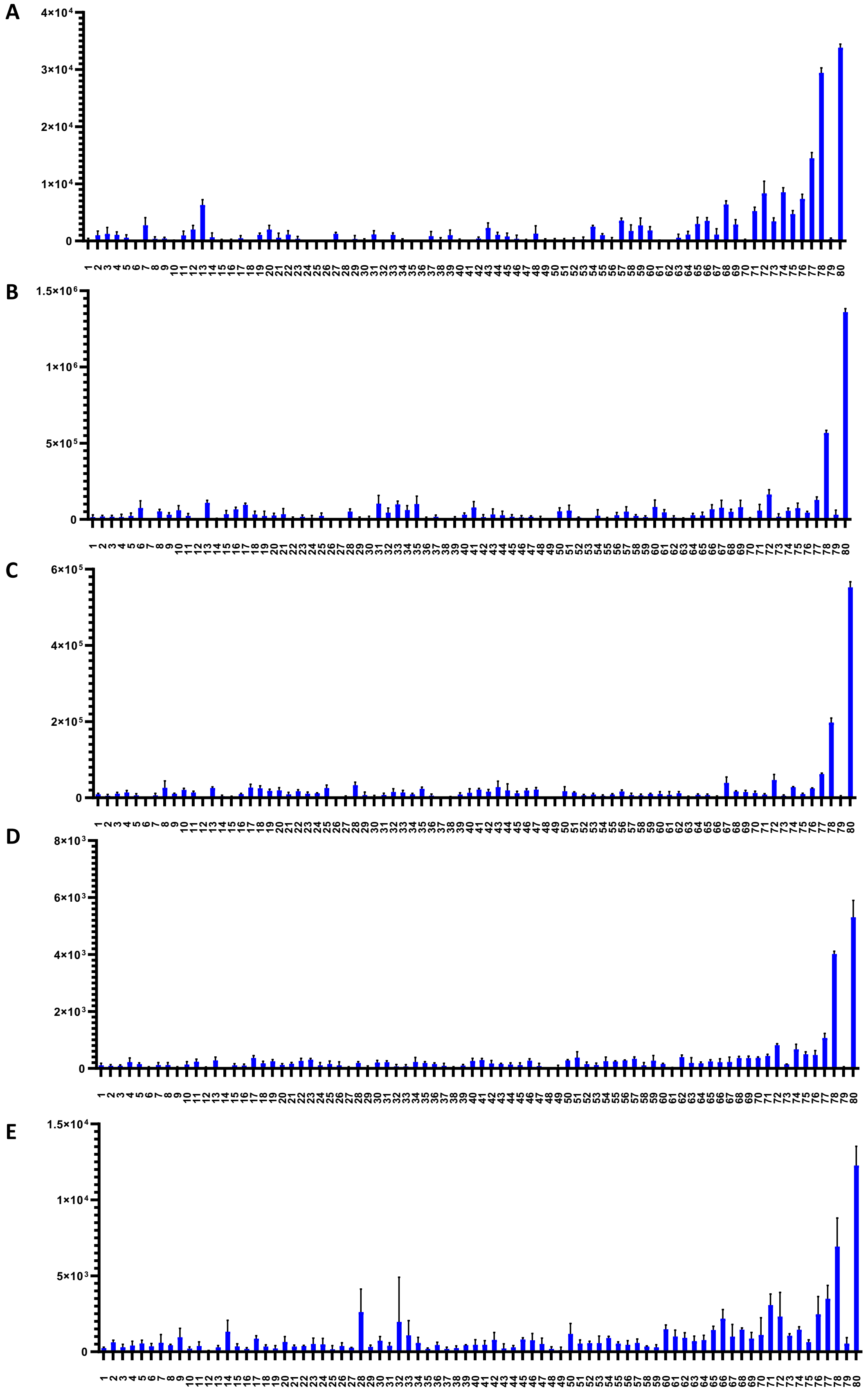

| | |
|---|---|
| **1** | IdoA-GlcNAc6S |
| **2** | GlcA-GlcNAc6S |
| **3** | IdoA2S-GlcNAc6S |
| **4** | IdoA2S-GlcNS6S |
| **5** | GlcA-GlcNAc-GlcA-GlcNAc |
| **6** | GlcA-GlcNAc-IdoA-GlcNAc |
| **7** | GlcA-GlcNAc-IdoA2S-GlcNAc |
| **8** | GlcA-GlcNAc-GlcA2S-GlcNAc |
| **9** | GlcA-GlcNAc-IdoA-GlcNAc6S |
| **10** | IdoA-GlcNAc6S-GlcA-GlcNAc |
| **11** | IdoA2S-GlcNAc-GlcA-GlcNAc |
| **12** | IdoA-GlcNAc-IdoA-GlcNAc6S |
| **13** | IdoA-GlcNAc6S-IdoA-GlcNAc |
| **14** | GlcA-GalNAc-GlcA-GalNAc4S |
| **15** | IdoA-GlcNS-IdoA-GlcNAc |
| **16** | IdoA-GlcNAc-IdoA-GlcNS |
| **17** | IdoA-GlcNAc6S-IdoA-GlcNAc6S |
| **18** | IdoA-GlcNAc6S-GlcA-GlcNAc6S |
| **19** | GlcA-GlcNAc6S-IdoA-GlcNAc6S |
| **20** | GlcA-GlcNAc6S-GlcA-GlcNAc6S |
| **21** | GlcA-GlcNAc-GlcA2S-GlcNAc6S |
| **22** | GlcA-GlcNAc-IdoA-GlcNS6S |
| **23** | GlcA-GlcNAc-IdoA2S-GlcNAc6S |
| **24** | IdoA2S-GlcNAc6S-GlcA-GlcNAc |
| **25** | GlcA-GalNAc4S-GlcA-GalNAc4S |
| **26** | GlcA-GlcNS6S-GlcA-GlcNAc |
| **27** | IdoA-GlcNS-IdoA-GlcNS |
| **28** | GlcA-GlcNAc6S-IdoA2S-GlcNAc6S |
| **29** | IdoA-GlcNAc6S-IdoA2S-GlcNAc6S |
| **30** | GlcA-GlcNS-IdoA2S-GlcNS |
| **31** | GlcA-GlcNS-GlcA2S-GlcNS |
| **32** | GlcA-GlcNAc-IdoA2S-GlcNS6S |
| **33** | GlcA-GlcNS-IdoA-GlcNS6S |
| **34** | IdoA-GlcNS6S-GlcA-GlcNS |
| **35** | IdoA2S-GlcNAc6S-GlcA-GlcNAc6S |
| **36** | IdoA2S-GlcNS-GlcA-GlcNS |
| **37** | GlcA-GlcNS6S-IdoA-GlcNS |
| **38** | IdoA-GlcNS-IdoA-GlcNS6S |
| **39** | GlcA-GlcNAc6S-GlcA2S-GlcNAc6S |
| **40** | IdoA-GlcNS6S-GlcA-GlcNS6S |
| **41** | GlcA-GlcNS6S-IdoA-GlcNS6S |
| **42** | GlcA-GlcNS6S-GlcA-GlcNS6S |
| **43** | IdoA-GlcNS6S-IdoA-GlcNS6S |
| **44** | GlcA-GlcNS-GlcA2S-GlcNS6S |
| **45** | GlcA-GlcNS-IdoA2S-GlcNS6S |
| **46** | IdoA2S-GlcNS6S-GlcA-GlcNS |
| **47** | IdoA2S-GlcNAc6S-IdoA2S-GlcNAc6S |
| **48** | IdoA-GlcNS-IdoA2S-GlcNS6S |
| **49** | IdoA2S-GlcNS6S-IdoA-GlcNAc6S |
| **50** | GlcA-GlcNS6S-IdoA2S-GlcNS6S |
| **51** | IdoA2S-GlcNS6S-GlcA-GlcNS6S |
| **52** | IdoA-GlcNS6S-IdoA2S-GlcNS6S |
| **53** | GlcA-GlcNS3S-IdoA2S-GlcNS6S |
| **54** | IdoA2S-GlcNS6S-IdoA2S-GlcNS |
| **55** | GlcA-GlcNS6S-GlcA2S-GlcNS6S |
| **56** | IdoA2S-GlcNS6S-IdoA2S-GlcNS6S |
| **57** | GlcA-GlcNS3S6S-IdoA2S-GlcNS6S |
| **58** | GlcA-GlcNAc-IdoA2S-GlcNAc6S-GlcA-GlcNAc |
| **59** | GlcA-GlcNS-IdoA2S-GlcNS6S-GlcA-GlcNS |
| **60** | GlcA-GlcNAc-IdoA2S-GlcNS6S-IdoA2S-GlcNS |
| **61** | GlcA-GlcNS6S-GlcA-GlcNS6S-GlcA-GlcNS6S |
| **62** | GlcA-GlcNS6S-IdoA-GlcNS6S-GlcA-GlcNS6S |
| **63** | GlcA-GlcNS6S-GlcA-GlcNS6S-IdoA-GlcNS6S |
| **64** | GlcA-GlcNS6S-IdoA-GlcNS6S-IdoA-GlcNS6S |
| **65** | GlcA-GlcNS-IdoA2S-GlcNS6S-IdoA2S-GlcNS |
| **66** | GlcA-GlcNAc6S-IdoA2S-GlcNS6S-IdoA2S-GlcNS |
| **67** | GlcA-GlcNS6S-IdoA2S-GlcNS6S-GlcA-GlcNS6S |
| **68** | GlcA-GlcNS6S-IdoA2S-GlcNS6S-IdoA-GlcNS6S |
| **69** | GlcA-GlcNS6S-GlcA-GlcNS6S-IdoA2S-GlcNS6S |
| **70** | GlcA-GlcNS6S-IdoA-GlcNS6S-IdoA2S-GlcNS6S |
| **71** | GlcA-GlcNS6S-IdoA2S-GlcNS6S-IdoA2S-GlcNS |
| **72** | GlcA-GlcNS6S-IdoA2S-GlcNS3S6S-GlcA-GlcNS6S |
| **73** | GlcA-GlcNS6S-IdoA2S-GlcNS3S6S-IdoA-GlcNS6S |
| **74** | GlcA-GlcNS6S-GlcA-GlcNS3S6S-IdoA2S-GlcNS6S |
| **75** | GlcA-GlcNS6S-IdoA-GlcNS3S6S-IdoA2S-GlcNS6S |
| **76** | GlcA-GlcNS6S-IdoA2S-GlcNS6S-IdoA2S-GlcNS6S |
| **77** | GlcA-GlcNS6S-IdoA2S-GlcNS3S6S-IdoA2S-GlcNS6S |
| **78** | IdoA2S-GlcNS6S-IdoA2S-GlcNS6S-IdoA2S-GlcNS6S |
| **79** | GlcA-GlcNS6S-IdoA-GlcNS-IdoA2S-GlcNS6S-IdoA-GlcNAc6S |
| **80** | IdoA2S-GlcNS6S-IdoA2S-GlcNS6S-IdoA2S-GlcNS6S-IdoA2S-GlcNS6S |

**Figure S1.** Binding of synthetic heparan sulfate oligosaccharides to SARS-CoV-2-related proteins, including (A) SARS-CoV-2 RBD-His (100 µg/mL), (B) SARS-CoV-2 RBD-Fc (10 µg/mL), (C) SARS-CoV-2 RBD trimer-streptavidin (3 µg/mL), (D) SARS-CoV-2 spike monomer-His (3 µg/mL), and (E) SARS-CoV-2 spike trimer-streptavidin (10 µg/mL). (F) Compounds numbering and structures of the heparan sulfate library.

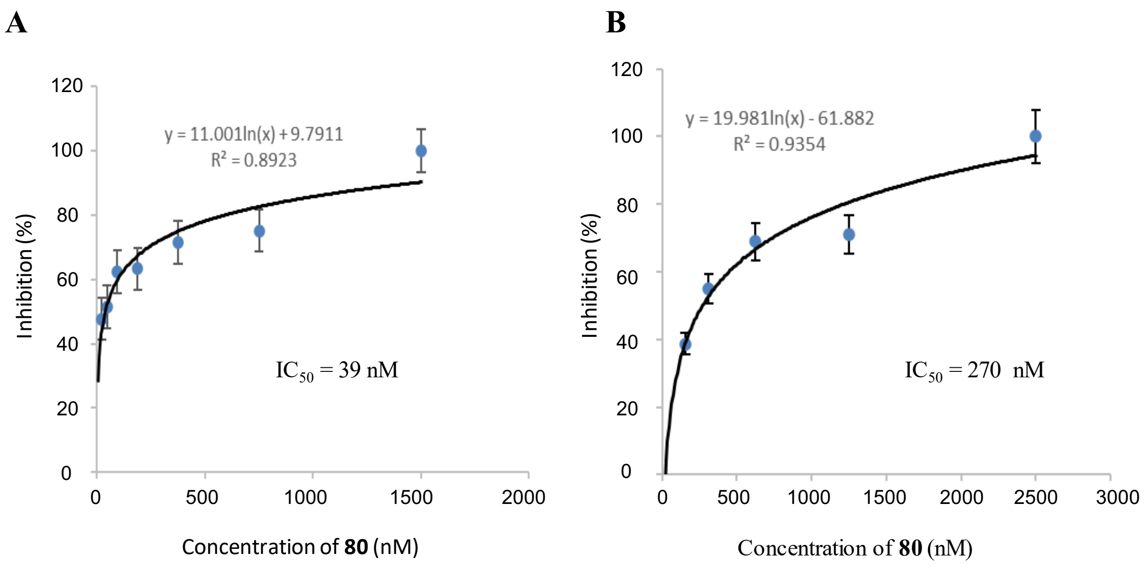


**Figure S2.** Inhibition of the binding of (A) SARS-CoV-2 spike (150 nM) and (B) RBD (5 μM) to HG-Hep by compound **80** measured by SPR.

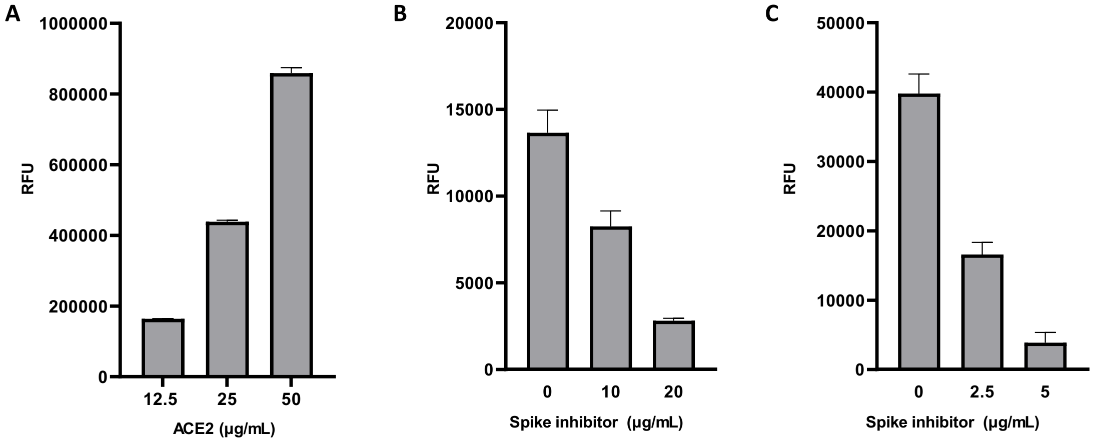


**Figure S3.** (A) Dose response of ACE2-Fc-biotin printed on streptavidin-coated slides at the indicated concentrations and detected by a goat-anti-human Fc antibody conjugated with AlexaFluor 647. (B) Humans ACE2 efficiently inhibits RBD-ACE2 and (C) spike-ACE2 binding. In (B) and (C) ACE2-Fc-biotin was printed at 50 µg/mL.

**References**

1. Stadlbauer, D.; Amanat, F.; Chromikova, V.; Jiang, K.; Strohmeier, S.; Arunkumar, G. A.; Tan, J.; Bhavsar, D.; Capuano, C.; Kirkpatrick, E.; Meade, P.; Brito, R. N.; Teo, C.; McMahon, M.; Simon, V.; Krammer, F. SARS-CoV-2 seroconversion in humans: A detailed protocol for a serological assay, antigen production, and test setup. *Curr. Protoc. Microbiol.* **2020**, *57* (1), e100.

2. Amanat, F.; Stadlbauer, D.; Strohmeier, S.; Nguyen, T. H. O.; Chromikova, V.; McMahon, M.; Jiang, K.; Arunkumar, G. A.; Jurczyszak, D.; Polanco, J.; Bermudez-Gonzalez, M.; Kleiner, G.; Aydillo, T.; Miorin, L.; Fierer, D. S.; Lugo, L. A.; Kojic, E. M.; Stoever, J.; Liu, S. T. H.; Cunningham-Rundles, C.; Felgner, P. L.; Moran, T.; Garcia-Sastre, A.; Caplivski, D.; Cheng, A. C.; Kedzierska, K.; Vapalahti, O.; Hepojoki, J. M.; Simon, V.; Krammer, F. A serological assay to detect SARS-CoV-2 seroconversion in humans. *Nat. Med.* **2020**, *26* (7), 1033-1036.

3. Bouwman, K. M.; Tomris, I.; Turner, H. L.; van der Woude, R.; Bosman, G. P.; Rockx, B.; Herfst, S.; Haagmans, B. L.; Ward, A. B.; Boons, G. J.; de Vries, R. P., Multimerization- and glycosylation-dependent receptor binding of SARS-CoV-2 spike proteins. *bioRxiv* **2020**, Preprint (DOI: 0.1101/2020.09.04.282558).

4. Zhang, F.; Zheng, L.; Cheng, S.; Peng, Y.; Fu, L.; Zhang, X.; Linhardt, R. J., Comparison of the interactions of different growth factors and glycosaminoglycans. *Molecules* **2019,** *24* (18), 3360.

5. Sun, Y.; Vandenbriele, C.; Kauskot, A.; Verhamme, P.; Hoylaerts, M. F.; Wright, G. J., A human platelet receptor protein microarray identifies the high affinity immunoglobulin E receptor subunit α (FcεR1α) as an activating platelet endothelium aggregation receptor 1 (PEAR1) ligand. *Mol. Cell. Proteomics* **2015,** *14* (5), 1265-1274.
